# A familial case of Syndactyly type IV due to a novel duplication of ~222.23 kb covering exons 2-17 of the LMBR1 gene: a case report

**DOI:** 10.1101/420588

**Authors:** Lijing Shi, Hui Huang, Qiuxia Jiang, Rongsen Huang, Wanyu Fu, Liangwei Mao, Xiaoming Wei, Huanhuan Cui, Keke Lin, Licheng Cai, You Yang, Yuanbai Wang, Jing Wu

**Affiliations:** Quanzhou Women’s and Children’s Hospital, 700 Fengze Street, Quanzhou, Fujian, 362000, China; BGI Genomics, BGI-Shenzhen, Shenzhen 518083, China; BGI-Wuhan, BGI-Shenzhen, Wuhan, 430074, China; BGI-Guangzhou Medical Laboratory, BGI-Shenzhen, Guangzhou 510006, China

**Keywords:** Syndactyly type IV, *LMBR1*, triphalangeal thumb-polysyndactyly syndrome, tibial and fibulal shortening, next generation sequencing.

## Abstract

Syndactyly is one of the most frequent hereditary limb malformations with clinical and genetical complexity. Autosomal dominant Syndactyly type IV (SD4) is a very rare form of syndactyly, occurring as a result of heterozygous mutation in an SHH regulatory element (ZRS) that resides in intron 5 of the *LMBR1* gene on chromosome 7q36.3. The SD4 is characterized by complete cutaneous syndactyly of all fingers, cup-shaped hands due to flexion of the fingers and accompanied by polydactyly. Here, we firstly reported a big Chinese family, manifesting cup-shaped hands consistent with SD4 and intrafamilial heterogeneity in clinical phenotype of tibial and fibulal shortening, triphalangeal thumb-polysyndactyly syndrome (TPTPS). Genetically, we identified a novel duplication of ∼222.23 kb covering exons 2-17 of the *LMBR1* gene in this family by next generation sequencing. This case expands our new clinical understanding of SD4 phenotype.

## INTRODUCTION

Syndactyly is one of the most frequent hereditary limb malformations(Tonkin, 2009). The occurrence rate of Syndactyly is variable due to geographical and registry differences, ranging from 1.1/10,000 in the northern Netherlands(Vasluian et al., 2013) to 1.3/10,000 in New York State (Goldfarb, Shaw, Steffen, & Wall, 2017). The incidence of syndactyly in China is **7.4**/10,000 in 1998-2009 (Sun, Xu, Liang, Li, & Tang, 2011). Currently eight pathogenic genes for syndactyly have been found, which are relative with nonsyndromic syndactyly types I-c, II-a, II-b, III, IV, V, VII, VIII (Ahmed, Akbari, Emami, & Akbari, 2017). Syndactyly type IV (SD4; OMIM 186200), also known as Haas polysyndactyly, is an autosomal dominant condition occurring as a result of heterozygous mutation in an SHH regulatory element (ZRS) that resides in intron 5 of the *LMBR1* gene (the limb development membrane protein 1 gene)on chromosome 7q36.3. The SD4 is characterized by complete cutaneous syndactyly of all fingers, cup-shaped hands due to flexion of the fingers, accompanied by polydactyly. The phenotypic and the genotypic variability in syndactyly may pose a diagnostic challenge for clinicians. Here, we reported a big Chinese family who have a novel duplication of ∼222.23 kb covering exons 2-17 of the *LMBR1* gene, manifest cup-shaped hands consistent with SD4 and intrafamilial heterogeneity in clinical phenotype of tibial and fibulal shortening, triphalangeal thumb-polysyndactyly syndrome (TPTPS) by next generation sequencing.

## CLINICAL REPORT

The proband (II-6) (Fig. 1A) was a 26-year-old woman with twin pregnancy, presented malformed hands and fingers. Bilateral hands are complete syndactyly and six toes of the left foot was preaxial polydactyly, syndactyly of the right foot was between /4/5 toes) When she was at 22 weeks pregnant, prenatal ultrasound was performed in our hospital. B-ultrasound revealed that one of the twins had fetal hands in a fist-shaped posture and there was no finger-to-finger interval, and hands and fingers deformity were not excluded. We further found she had an abnormal pregnancy-Labor history. Her first fetus was aborted due to the same abnormal ultrasound result during 20 weeks of pregnancy. For investigating her family history, we learned that she was from a big Chinese family segregating autosomal dominant non-syndromic Syndactyly in Fujian Province. 8 members of three generations are affected with continuous transmission, including 3 males and 5 females (Figure 1A). Two of these cases (III-1, III-4) were syndactyly confirmed by prenatal ultrasonography and finally induced labor. None of his family members exhibited intellectual anomalies.

**Figure 1.**
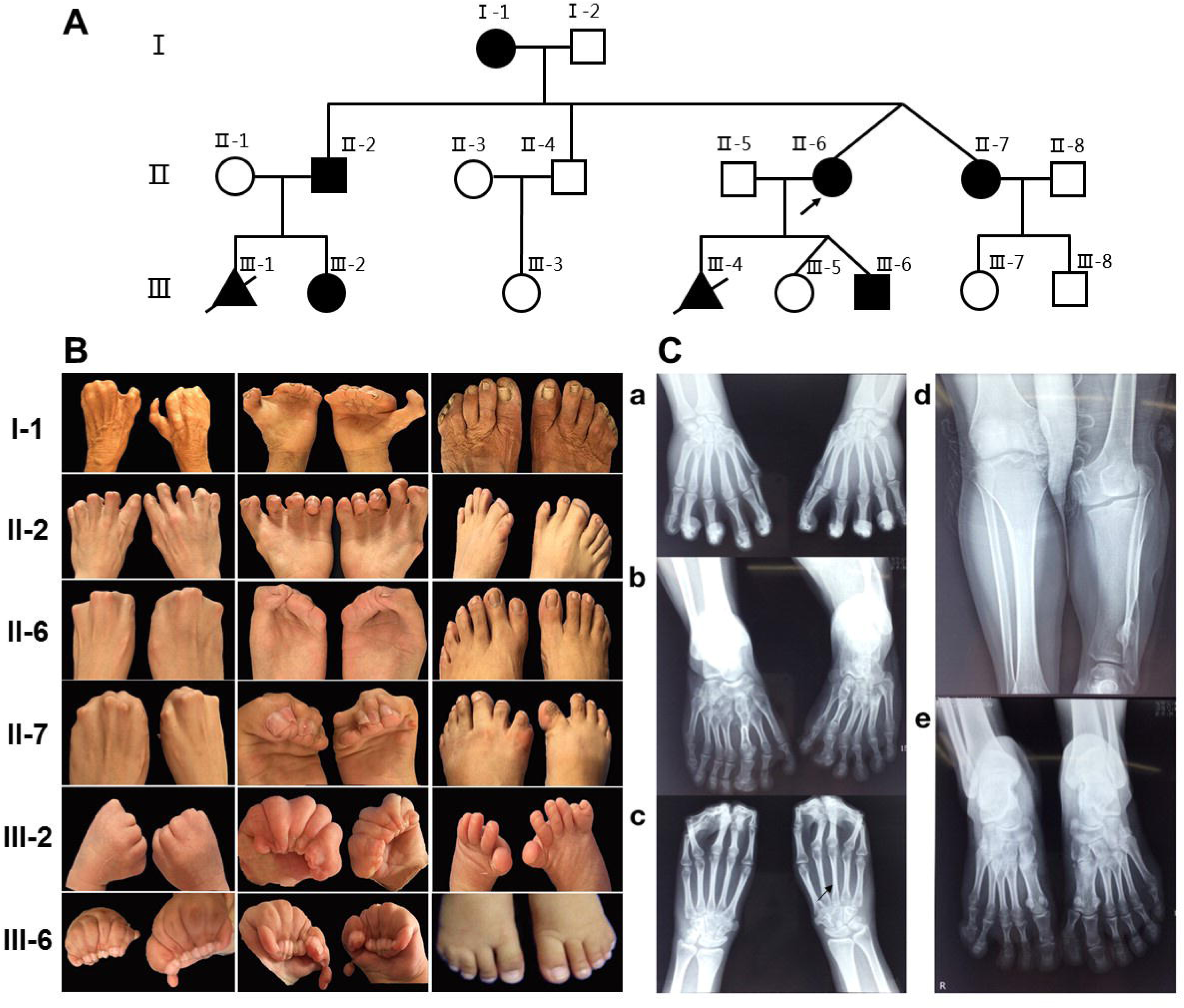
A Pedigree of the 3 generation family with Syndactyly. Family members with Syndactyly are marked with shading. Individuals labeled with a triangles and slash were abortion. Arrow indicates the proband (II-6). **B Images of limb features in the family. C X-ray images of hands and feet PA in II-2 and the proband (II-6).** a) Hands PA of II2, postoperation: An extra metacarpal and two phalanxes are observed at the lateral part of the first metacarpal on the left hand, and the soft tissue of the extra metacarpal and that of thumb (left hand) is fused. Another extra metacarpal and one phalanx is observed at the lateral part of the first metacarpal on the right hand, and the soft tissue of the extra metacarpal and that of thumb (right hand) is fused. The two hands of the distal phalangeal flexion overlap. b) Feet PA of II2: Both feet are varus and the tarsal bones are markedly shortened. Six tarsal and phalanx bones are found on left feet, and seven tarsal and phalanx bones are found on the right feet; Density reduction occurs in the marrow cavities of the left second tarsal bone and the right third tarsal bone; Bone sclerosis is detected in surrounding part; second left and third right proximal phalanx thickens; second left and third right distal phalanx thickens and deforms significantly. d) Bilateral tibial and fibular PA & LAT of II2: The left side of the tibia is markedly shortened, the density of the medullary cavity in the distal part of the left fibula decreases, and the surrounding bone shows sclerosis. There is a bony eminence in the local cortex. c) Hands PA of II-6: The middle and the distal phalanx overlap, and the soft tissue of the distal part of both hands is fused. e) Feet PA of II-6:A polydactylism is observed at the lateral part of the hallux on left foot, which contains two sections of short phalange with the formation of the joint between phalanges, proximal phalanx bones and the first tarsal joint, the size and morphology of the first tarsal bone is normal. The distal phalanx of the fifth toes on left foot enlarges and deforms.

By a physical examination, we found the clinical phenotype of family members is similar but varied. The presentation in most of the affected patients was bilateral and symmetrical, accompanied by syndactyly and polydactyly (Figure 1B). Phenotypic spectrum in all the affected subjects can be seen supplementary material table 1. Some affected individuals have other abnormalities in their fingers and lower limb. In appearance, I-1 presented with triphalangeal thumb on her right hand, and II-2, who had hand surgical operation, appeared limply walking.

Clinical X-ray examination of the proband (II-6) (Figure 1-c,e) and II-2 (Figure 1C-a,b,d) revealed that the characteristics of the deformity of both hands and feet and the changes of the bilateral tibial and fibular. II-2 represented by unilateral shortened tibia and fibula (Figure 1C-d), triphalangeal thumb on his right hand (Figure 1C-c). X-rays from other family members were not available.

## MOLECULAR ANALYSES

Our research observed the tenets of the Helsinki Declaration and was approved by the institutional review board of Quanzhou Woman’s and children’s Hospital. Six members of a Han Chinese non-consanguineous family were recruited. Written consent for reporting clinical results was obtained from all the participants. Genomic DNA was extracted from patients’ peripheral blood samples and were sequenced using a BGISEQ-500 sequencer, which average coverage depth is nearly 300×. The targeted sequences were captured using the Genetic disease chip, which contained 3299 genes and covered 6 genes (*HOXD13, FBLN1, ZRS(LMBR1), LRP4, FGF16, GREM1-FMN1)* related to Syndactyly according to OMIM. Variants analysis and filtering was performed as previously reported(Patch et al., 2018; Zheng et al., 2018). The identified variant was confirmed by Quantitative PCR (qPCR). The breakpoint mapping and size of duplication mutation (156,460,343-156,682,575,222Kb) were confirmed in the proband by low-coverage pair-end whole-genome sequencing as previously reported (Dan et al., 2012) (supplementary materials figure1)

Targeted gene testing of six patients (II-6, III-6, II-7, III-7, II-4, II-2) were performed to make molecular pathogenic diagnosis. Duplication of Exon2-Exon17 in the gene coding region of *LMBR1* (NM_022458. 3) was detected, and four patients (II-6, III-6, II-7, II-2)both carried the heterozygous mutation, however two unaffected family members (II-4, III-7) have no the duplication mutation. Quantitative PCR (qPCR) for pedigree analysis confirmed the mutation is co-segregation in this family (Figure 2).

**Figure 2.**
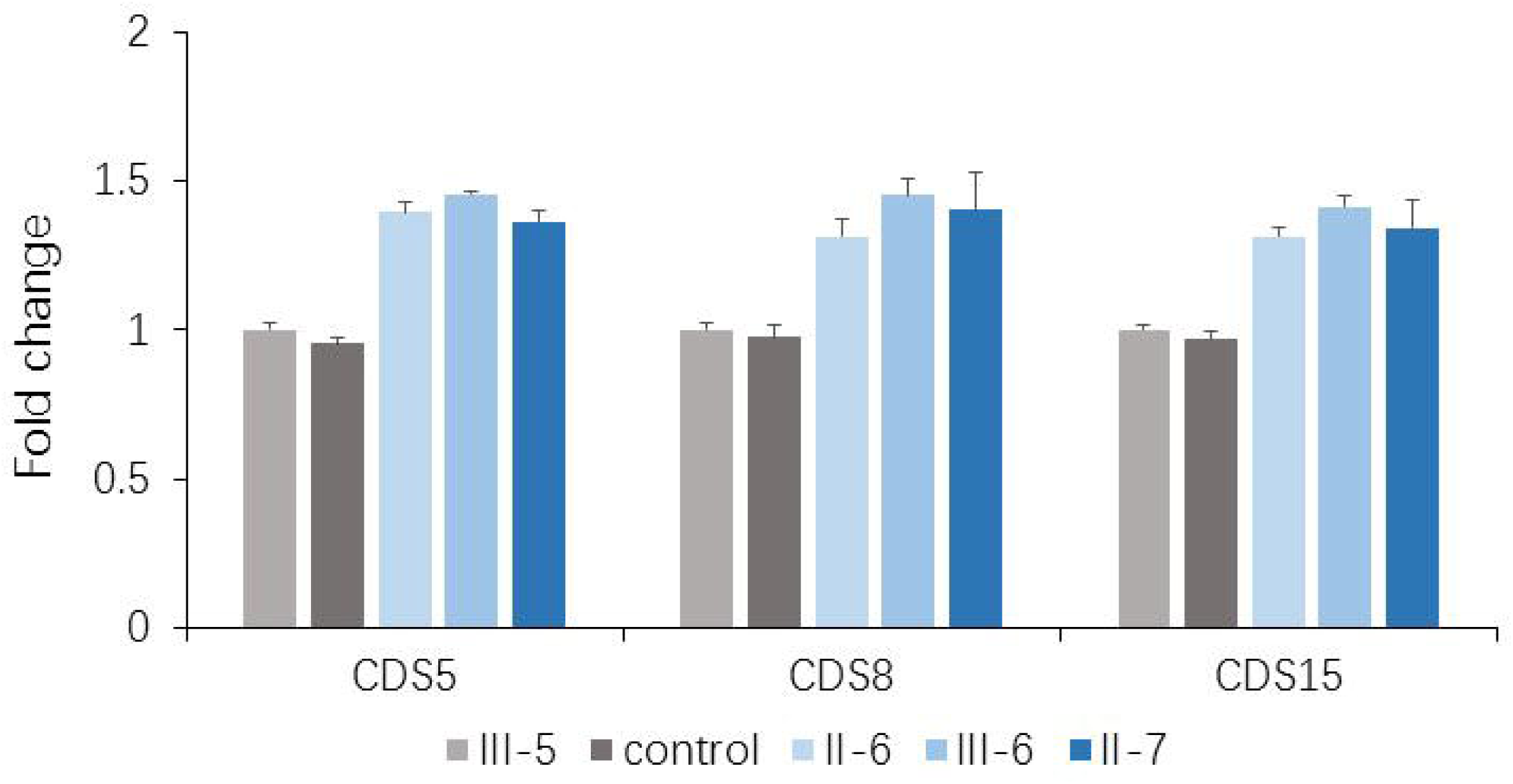
Quantitative real-time PCR validation of *LMBR1* duplication. Measurement was performed at three different positions (CDS5, CDS8 and CDS15) and normalized to the ACTB gene.

## DISCUSSION

Here we firstly reported the clinical manifestations and molecular features of a big Chinese family with autosomal dominant non-syndromic SD4 in Fujian Province. By targeted gene testing and qPCR analysis, we found all the affected patients had a 222.23 kb duplication of exons 2-17 in *LMBR1* gene which is absent in unaffected family members. Reported studies found that genomic large duplications in the ZRS region of *LMBR1*, covering approximately from 73-kb to 589-kb were responsible for human SD4 and TPTPS in many different country families (Klopocki et al., 2008; Wieczorek et al., 2010). The ZRS locates approximately 1 Mb away from SHH (Lettice et al., 2003), which controls the expression of SHH in the developing limb, and is conserved among mammals and fish (Lettice et al., 2003). The Lmbr1 mutant mouse displayed syndactyly involving digits II to V (Al-Qattan, Shamseldin, Al Mazyad, Al Deghaither, & Alkuraya, 2013), which is in line with the phenotypes of SD4.

In this family, all patients had the typical clinical phenotype of SD4, Cup-shaped hands caused of the fingers together with cutaneous syndactyly, companied with limited flexion of the fingers, but no fusion of phalanges and metacarpus. Meanwhile, the clinical phenotypic diversity of our family members is characterized by varying degrees of syndactyly and polydactyly in different fingers, skin fusion, pre-axial polydactyly and fused fingernails. The presentation in most of the affected patients was bilateral and symmetrical (Figure 1B). Special attention should be paid to differences in the malformations of the twins. The right toe of II-7 is normal and the left toe of II-7 is an extra Postaxial toe. The right fourth and fifth toes of II-6 are syndactyly and the left toe of II-6 are two extra Postaxial toes. The family also had a complex limb phenotype characterized by triphalangeal thumb (I-1, II-2), pre-axial polydactyly, extra preaxial toe. In addition, III-6 had a congenital cleft lip and palate, which has been treated surgically. The phenotype of II-2 is the most variable. His hands are complete syndactyly (surgically treated) and triphalangeal thumb, with seven-toes deformity and the asymmetrical syndactyly on the left and right toes and accompanied by 1-2 small toes of both feet, the third big toe shaped like a thumb.

Patients with duplications involving ZRS are mainly diagnosed of TPTPS and /or SD4 from reported families with 11 duplications identified (Dai et al., 2013), its clinical phenotype had almost no lower limb development. In these cases, only two cases reported SD4 complicated with fibula dysplasia, but relevant data were insufficient or absent for the X image of fibula dysplasia (Wu et al., 2009) or the duplication fragment size (Sato et al., 2007). In comparison, II-2 in our case was identified markedly shortened tibia and fibula on his left leg, and triphalangeal thumb on his right hand but it is not sure on his left hand because of all bent and overlapping fingers by X ray. So the duplication of exons 2-17 in *LMBR1* gene may cause SD4 and TPTPS with shortened tibia and fibula. SD4 and TPTPS have been reportedly considered as a continuum of phenotypes(Dai et al., 2013), but phenotype of limb malformation were found only in two reported patients with varying degrees of severity, and did not appear to be directly related to the size of the repeated duplication fragments, such as 589 kb duplication related with TPTPS (Klopocki et al., 2008) completely cover our duplication region, 115 kb duplication related with TPTPS and SD4, PAP of toe was contained in our region (Dai et al., 2013) suggesting that other complex mechanisms might lead to related phenotypes or clinical heterogeneity to further interpretation.

In summary, we firstly described an affected family with heterozygous duplication of exons 2-17 in *LMBR1* gene who presented with SD4 accompanied with a diverse phenotype of triphalangeal thumb, and tibia and fibula shortening in Quanzhou City, Fujian Province. It again confirmed the genetic homogeneity of TPTPS and SD4, and demonstrated intrafamilial phenotypic heterogeneity. It also suggests that in future genetic counseling, we should strengthen the popularization of education to the disease so as to guide reproduction in this area.

## ACKNOWLEDGEMENT

We thank sincerely for the participation of all members from this Chinese family.

## DISCLOSURE OF INTERESTS

None

## CONFLICT OF INTERESTS

The authors have declared no conflicting interests.

